# Characterization of a new cytorhabdovirus discovered in papaya (*Carica papaya*) plantings of Ecuador and its relationship with a bean-infecting strain from Brazil

**DOI:** 10.1101/605139

**Authors:** Andres X. Medina-Salguero, Juan F. Cornejo-Franco, Samuel Grinstead, Dimitre Mollov, Joseph D. Mowery, Francisco Flores, Diego F. Quito-Avila

## Abstract

The complete genome of a new rhabdovirus infecting papaya (*Carica papaya* L.) was sequenced and characterized. The genome consists of 13,469 nucleotides with six canonical open reading frames (ORFs) predicted from the antigenomic strand. In addition, two overlapping short ORFs were predicted between ORFs 3 and 4. Phylogenetic analyses using amino acid sequences from the nucleocapsid, glycoprotein and polymerase, grouped the virus with members of the genus *Cytorhabdovirus*, with rice stripe mosaic virus, yerba mate chlorosis-associated virus and Colocasia bobone disease-associated virus as closest relatives. The 3’ leader and 5’ trailer sequences were 144 and 167 nt long, respectively. Each end contains complementary sequences prone to form panhandle structures. The motif 3’-AUUCUUUUUG-5’, conserved across rhabdoviruses, was identified in all but one intergenic regions; whereas the motif 3’-ACAAAAACACA-5’ was found in three intergenic junctions. This is the first complete genome of a cytorhabdovirus infecting papaya. The virus was prevalent in commercial plantings of Los Ríos, the most important papaya producing province of Ecuador. During the final stage of this manuscript preparation, the genome of a bean-associated cytorhabdovirus became available. Nucleotide identity (97%) between both genomes indicated that the two viruses are strains of the same species, for which we propose the name papaya cytorhabdovirus E.

## Introduction

The *Rhabdoviridae*, a negative-sense RNA virus family, contains viruses that infect a wide range of hosts including vertebrates, invertebrates and plants [1]. Virions have a helical, bullet-shape morphology, surrounded by a host-derived membrane [2]. Rhabdovirus genomes range from 11 to 16 kilobases (kbp) with only non-segmented ones classically assigned to the taxa. However, virus species with bipartite genomes have recently been included in the family [3,4]. Based on host type, genomic organization and other biological features, rhabdoviruses currently are organized in 13 genera: *Cytorhabdovirus, Dichorhavirus, Ephemerovirus, Lyssavirus, Novirhabdovirus, Nucleorhabdovirus, Perhabdovirus, Sigmavirus, Sprivivirus, Tibrovirus, Tupoavirus, Varicosavirus* and *Vesiculovirus* (http:ictvonline.org/virusTaxonomy.asp). Next generation sequencing (NGS) techniques have led to the discovery of several novel rhabdovirus species for which new genera have been proposed [5,6,7].

The genome organization of rhabdoviruses has a canonical arrangement of five genes: 3’-N-P-M-G-L-5’ that encode the nucleocapsid protein, phosphoprotein, matrix protein, glycoprotein and the large polymerase, respectively. Terminal regions have non-coding regulatory sequences denoted, respectively, as 3’-leader (*l*) and 5’ trailer (*t*) [8-9]. Additional “accessory” genes have been observed in arrangements that differ among rhabdoviruses [10].

Plant-infecting rhabdoviruses have long been classified into either the *Cytorhabdovirus* or the *Nucleorhabdovirus* genus, based on their cytoplasmic or nuclear, respectively, site of accumulation in the cell [11,12]. This biological feature has been confirmed by phylogenetic relationships, which separate clearly the two groups. Recently, two new genera, *Dichorhavirus* and *Varicosavirus*, have been created to classify novel plant-infecting rhabdoviruses with bi-partite genomes [3-4]. Although more than 80 non-segmented plant rhabdoviruses have been reported, based on cytopathology studies, complete genomes are only available for a few members of each genus. A genomic feature common to both cyto- and nucleorhabdoviruses is the presence of an additional ORF between the P and M genes. The product of this ORF is considered the movement protein (MP) as it has been demonstrated to have cell-to-cell movement function [13,14]. Additional non-classical ORFs have been identified in the genomes of some plant rhabdoviruses, resulting in variations of the canonical genomic organization [15].

Plant infecting rhabdoviruses have been found in a wide range of hosts including monocots such as rice, maize, wheat and barley, and dicots such as potato, lettuce, carrot and strawberry, among others [12].

In papaya (*Carica papaya* L.), the occurrence of nucleorhabdoviruses in commercial plantings in Venezuela, Florida and Mexico was documented as early as 1980 [16,17,18]. In Venezuela, the virus was observed in tissue collected from trees showing a range of symptoms including leaf yellowing, apical necrosis and plant death [16]. In Florida, the virus was associated with droopy necrosis, a disorder that included bending of the upper section of the crown with a bunchy appearance which, at later stages, developed into necrosis and plant death [17]. However, no genomic sequences for papaya rhabdoviruses have been reported.

This study reports the complete genome sequence of a new cytorhabdovirus discovered from papaya plants in Ecuador, and provides genomic, phylogenetic and biological characterization of the virus.

## Materials and methods

### Virus source

In 2016, a papaya sentinel plant (cv. Sunrise), previously used as part of a study on the epidemiology of papaya virus Q (PpVQ) and its relationship with papaya ringspot virus (PRSV) [19,20], was maintained under greenhouse conditions for further investigation. The study was conducted in Los Ríos, the largest papaya producing province of Ecuador, where sentinel plants were scattered in a 2-year old field and monitored for four months. The selected plant was subjected to virus testing using the reverse-transcription (RT)-PCR assay described by Quito-Avila et al. [20]. The plant tested positive for PRSV but not for PpVQ. Additional viruses infecting this plant were further investigated using the approach described below.

### Sequencing and genome analyses

Total RNA was extracted from the selected plant using the protocol described by Quito-Avila, et al. [20]; followed by DNase treatment. Viral RNA was enriched by depleting plant rRNA. Preparation of Trueseq RNA library, followed by NGS on a HiSeq 4000 Illumina platform (100 paired-end reads) was performed at Macrogen (South Korea). Sequence reads were trimmed and assembled into contigs by CLC Workbench 11 (Qiagen USA). Contigs were analyzed using BLASTx from the *National Center for Biotechnology Information* (NCBI).

NGS data was verified by Sanger sequencing of RT-PCR amplified overlapping fragments, which were generated by specific primers. Terminal sequences were confirmed by RACE using total RNA as a template for cDNA, and specific primers near the ends as recommended by the manufacturer (Roche, Germany). Sequence comparisons, alignments and prediction of open reading frames (ORFs) were done using Geneious R 11, the NCBI conserved domain database [21] and the Swiss-model server [22]

### Transmission Electron Microscopy (TEM)

Leaf tissue was dissected into one mm pieces using a biopsy punch, fixed in 2% paraformaldehyde, 2.5% glutaraldehyde, 0.2% Tween-20, 0.5M Na cacodylate and processed in a Pelco BioWave microwave as previously described [23]. TEM grids were imaged at 80kV with a Hitachi HT-7700 transmission electron microscope (Hitachi High Tech America, Inc., Dallas, TX, USA).

### Phylogenetic analyses

Protein sequences corresponding to the nucleocapsid, glycoprotein and polymerase were downloaded from all the nucleo- and cytorhabdoviruses available in GenBank. In addition, cytorhabdovirus-like sequences annotated as part of the whitefly (*Bemisia tabaci*) genome (acc. numbers: KJ994255-KJ994264), were identified and included in the analysis. Amino acid sequences were aligned using structural information with Expresso [24]. The confidence of the multiple sequence alignments was measured with TCS[25], and unreliable alignment fragments were discarded. The best evolution model for each alignment was determined with MEGA7 [26] and used to build single gene and multi-locus phylogenies on BEAST v.1.8.4 [27]. Two chains of 1000000 MCMC were run for each protein and for the multi-locus concatenation. Convergence of the runs and effective sample size were observed in Tracer v.1.6. The two runs were combined with a 10% burn-in using LogCombiner and a consensus tree was built with TreeAnnotator [27].

### Virus detection and survey

Primers were designed to amplify a fragment of the viral nucleocapsid gene and used in a virus survey of commercial papaya plantings in five provinces (Los Ríos, Guayas, Manabí, Santa Elena and Sucumbíos) of Ecuador. Total RNA extraction and RT-PCR was done as described [20]. Up to five positive samples from each location were also used to amplify a fragment of the virus polymerase. Amplicons were cloned and sequenced, and comparisons determined the sequence variability based on geographic location. Multiple sequence alignments were performed using ClustalW [28].

## Results

### Sequencing

A total of 32,375,528 pair-end 100 nt reads were obtained from the NGS cDNA library. These reads were assembled into 61,033 contigs. More than 20 contigs were identified as associated with plant viruses. One contig >13 kb showed similarities to cytorhabdoviruses. The remaining contigs revealed similarity to PRSV. In subsequent analysis using Geneious R11 (Biomatters, New Zealand) these contigs were assembled into a ∼10 kb PRSV genome. Only about 75 thousand reads (0.23%) mapped to the cytorhabdovirus, while 3.8 million (11.68%) identified with the PRSV contig.

### Genome organization

The entire genome of the new virus (GenBank accession no. MH282832) is 13,469 nt. The antigenomic strand contains six major open reading frames (ORFs) arranged in a typical plant rhabdovirus organization (Fig 1). BLASTx searches performed on each ORF revealed homology to different members of the genus *Cytorhabdovirus.*

**Fig 1.**
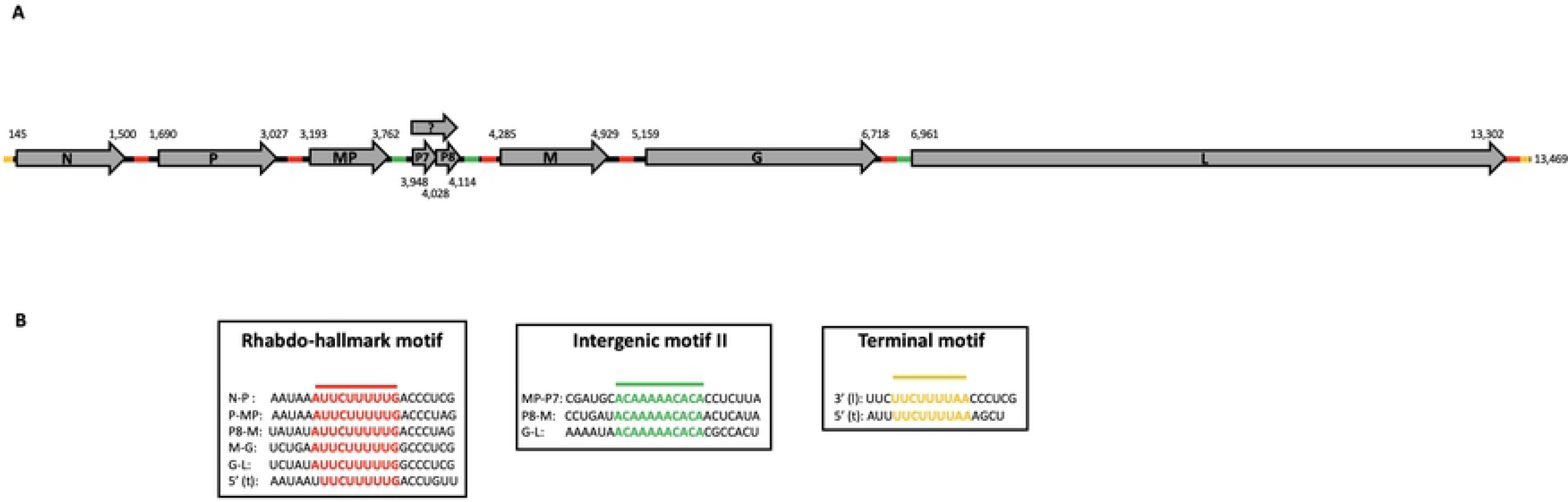
Genome organization of papaya rhabdovirus. A) Open reading frames (ORFs) with corresponding nucleotide positions are illustrated by the grey arrows, where N represents the nucleocapsid, P the phosphoprotein, MP the movement protein, M the matrix protein, G the glycoprotein and L the polymerase. P7 and P8 denote additional ORFs, which might be expressed as a fusion protein (denoted by the question mark). The presence of conserved motifs in intergenic or terminal regions is indicated by colored solid lines and detailed (3’ – 5’ orientation) in panel B.

ORF 1 (nt positions: 145 – 1,500) encodes the putative nucleocapsid protein (N), showing the highest identity (35%, amino acid level) with rice stripe mosaic virus (RSMV). ORF 2 (nt positions: 1,690 – 3,027) codes for a 445 aa protein with homology (15% aa identity) to the putative phosphoprotein (P) of yerba mate chlorosis associated virus (YMCaV) [29] and RSMV (13% identity).

ORF 3, (nt positions: 3,193 – 3,762), encodes a 189 aa protein with homology to the putative movement protein (MP) of YMCaV (23% aa identity).

ORF4, (nt positions: 4,285 – 4,929) encodes a 214 aa protein with 15% identity to the putative matrix protein (M) of Colocasia bobone disease-associated virus (CBDaV), a cytorhabdovirus recently discovered in Solomon Islands [30].

The theoretical product of ORF 5 (nt positions: 5,159 – 6,718) is a 519 aa protein homologous (20% aa identity) to the glycoprotein (G) of YmCaV, and the largest ORF 6 (nt positions: 6,961 – 13,302) encodes the putative polymerase (L) showing the highest aa identity to CBDaV (36%) and RSMV (33 %).

In addition to the six ORFs exhibiting the classical organization of cytorhabdoviruses, two short ORFs (here referred to as ORFs 7 and 8) were predicted, respectively, at nt positions 3,948 – 4,028 and 4,028 – 4,114 (Fig 1). Interestingly, ORFs 7 and 8 are arranged in a contiguous fashion with overlapping termination/initiation codons (e.g. <u>UA</u>**<u>A</u>UG**), typical of a reinitiation translation mechanism (RTM). RTM has been documented for several different negative-sense RNA viruses, including animal-infecting rhabdoviruses [31,5].

Several studies have shown that RTM occurs only after a very short ORF (less than 30 codons), which contains a termination upstream ribosome-binding site (TURBS) motif, located upstream the overlapping stop/start codons [32]. Despite fulfilling the short ORF condition (e.g. ORF 7 has 27 codons), TURBS-like motifs were not identified. However, the two contiguous ORFs are flanked by motifs (Fig 1), which are conserved across intergenic regions (see below), suggesting their legitimate expression.

Conserved domain database and pfam searches did not find homologues for ORFs 7 or 8. Nevertheless, a Swiss-model search, using the 54 aa polypeptide resulting from the hypothetical fusion of ORFs 7 and 8 as template, found a 28 aa-long hit (37.9 % identity) with the PRD1 bacteriophage minor capsid protein (RCSB accessions 1YQ5 and 1YQ8) [33].

### Intergenic regions

The new virus possesses intergenic regions with an average 36% GC content, except for the MP-P7 junction, which has 46.5%. The conserved motif 3’-AUUCUUUUUG-5’, a hallmark in rhabdoviruses [5], was found at each intergenic region, except for the one corresponding to the MP-P7 junction. Interestingly, a second motif (*intergenic motif II*), with the core sequence 3’-ACAAAAACACA-5’, was identified in junctions MP-P7, P8-M and G-L of the new virus (Fig 1). This motif was partially conserved in one or two intergenic junctions of other cyto- and nucleorhabdoviruses (Table 1).

**Table 1.**
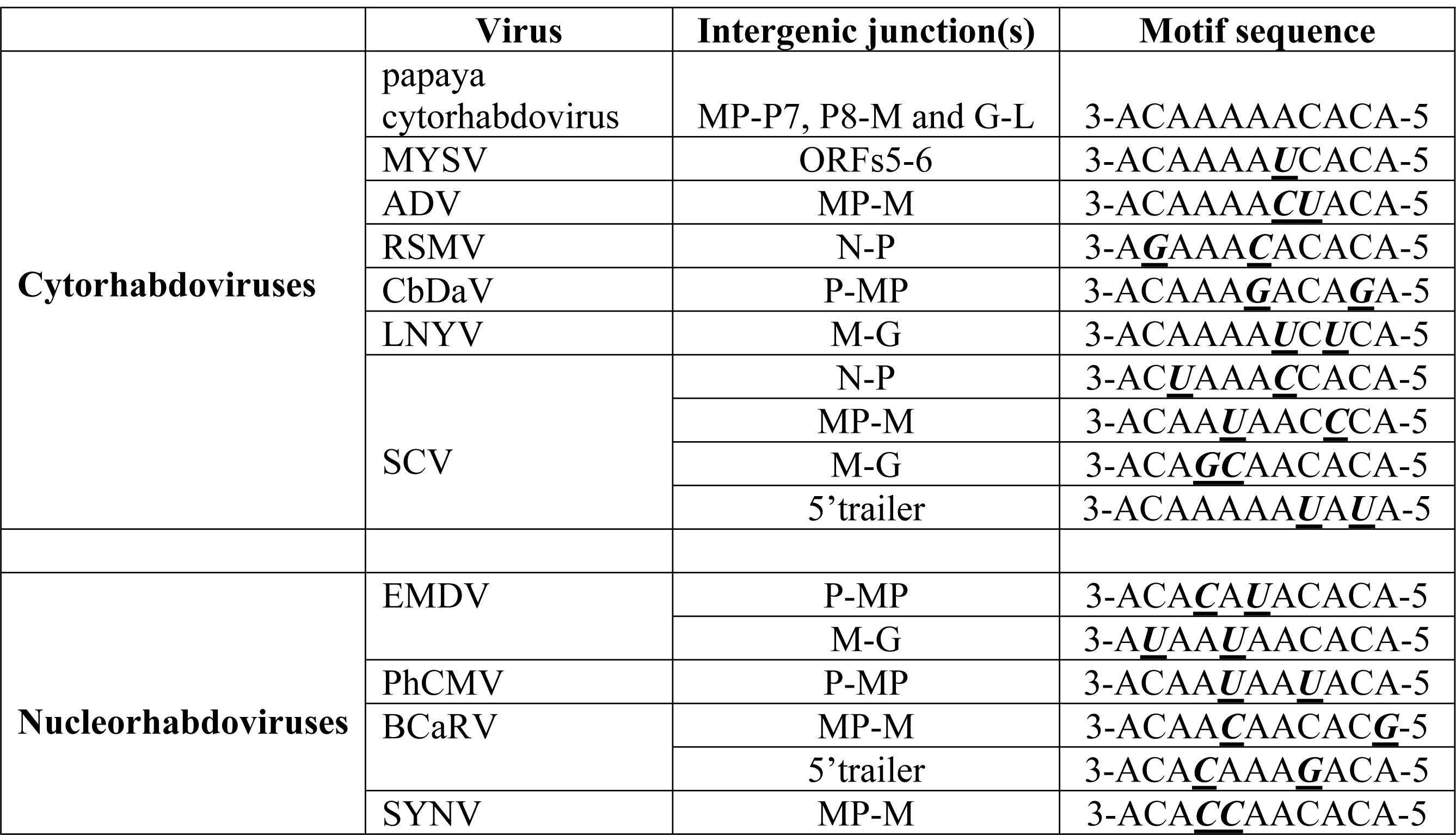
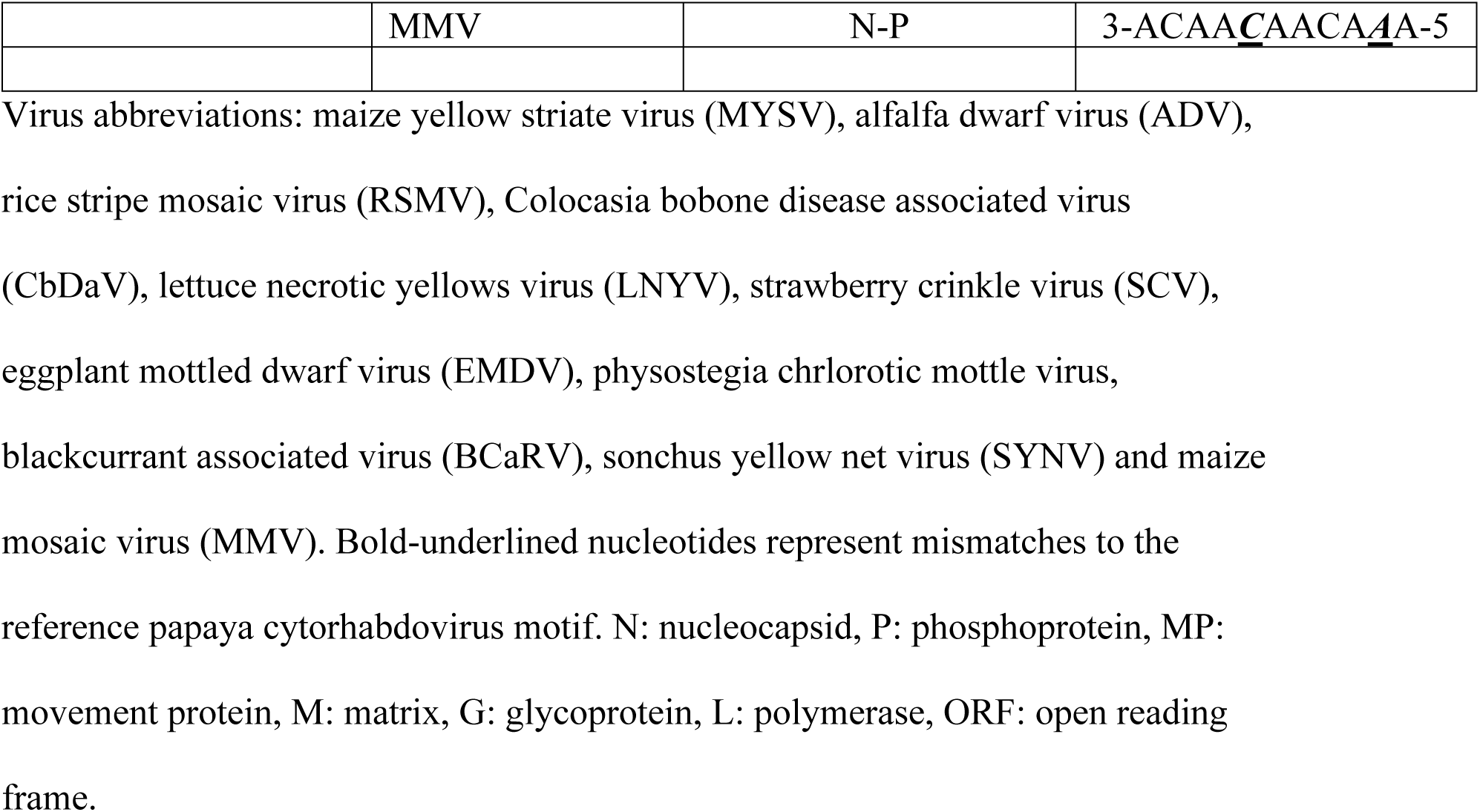
Presence of partially conserved intergenic motif II in cyto- and nucleorhabdoviruses.

A common feature found in the intergenic regions of the new virus was the presence of several complementary motifs with potential to form secondary structures. The complexity of hypothetical secondary structures varied across intergenic regions, with junctions N-P, M-G and G-L showing a greater number of complementary stretches S1 Fig.

### Terminal regions

The 3’ leader contains 144 nucleotides. This length is similar to those from other cytorhabdoviruses such as maize yellow striate virus (MYSV, 143 nt), NCMV (141 nt) and strawberry crinkle virus (SCV, 147 nt). Interestingly, the two most closely related cytorhabdoviruses, RSMV and YmCaV, have shorter 3’ leaders, with 89 nt and 73 nt, respectively. The 5’ trailer is 167 nt long, closely similar in size to its counterpart from YmCaV (140 nt).

Sequence examination of both terminal regions revealed the existence of four motifs with potential *cis*- or *trans*-acting interactions (Table 2). In addition, the rhabdovirus hallmark intergenic motif 3’-UUCUUUUUG-5’ was detected at the trailer region (nt positions 13,395-13,402) (Fig 1). Several additional motifs, with potential complementary interactions among each other, were identified in both terminal regions (Fig 2).

**Table 2.** Nucleotide motifs with potential *cis*- or *trans*-acting interactions in the terminal regions of the new papaya cytorhabdovirus.

| Motif | Genome position (nt) |  |
| --- | --- | --- |
|  | 3' leader | 5' trailer |
| 3'-GAUAAAA-5' | 12-18 |  |
| 3'-UUCUUUUAA-5' | 62-70 | 13,435-13,443 |
| 3'-ACUAUUUU-5' |  | 13,429-13,436 |
| 3'-AAAAUAGU-5' |  | 13,456-13,463 |
| 3'-UUCUUUUU-5' |  | 13,395-13,402 <sup>a</sup> |

Nucleotide (nt) positions with respect to the full genome length are indicated.
<sup>a</sup> This motif is identical to the rhabdovirus hallmark intergenic sequence.

**Fig 2.**
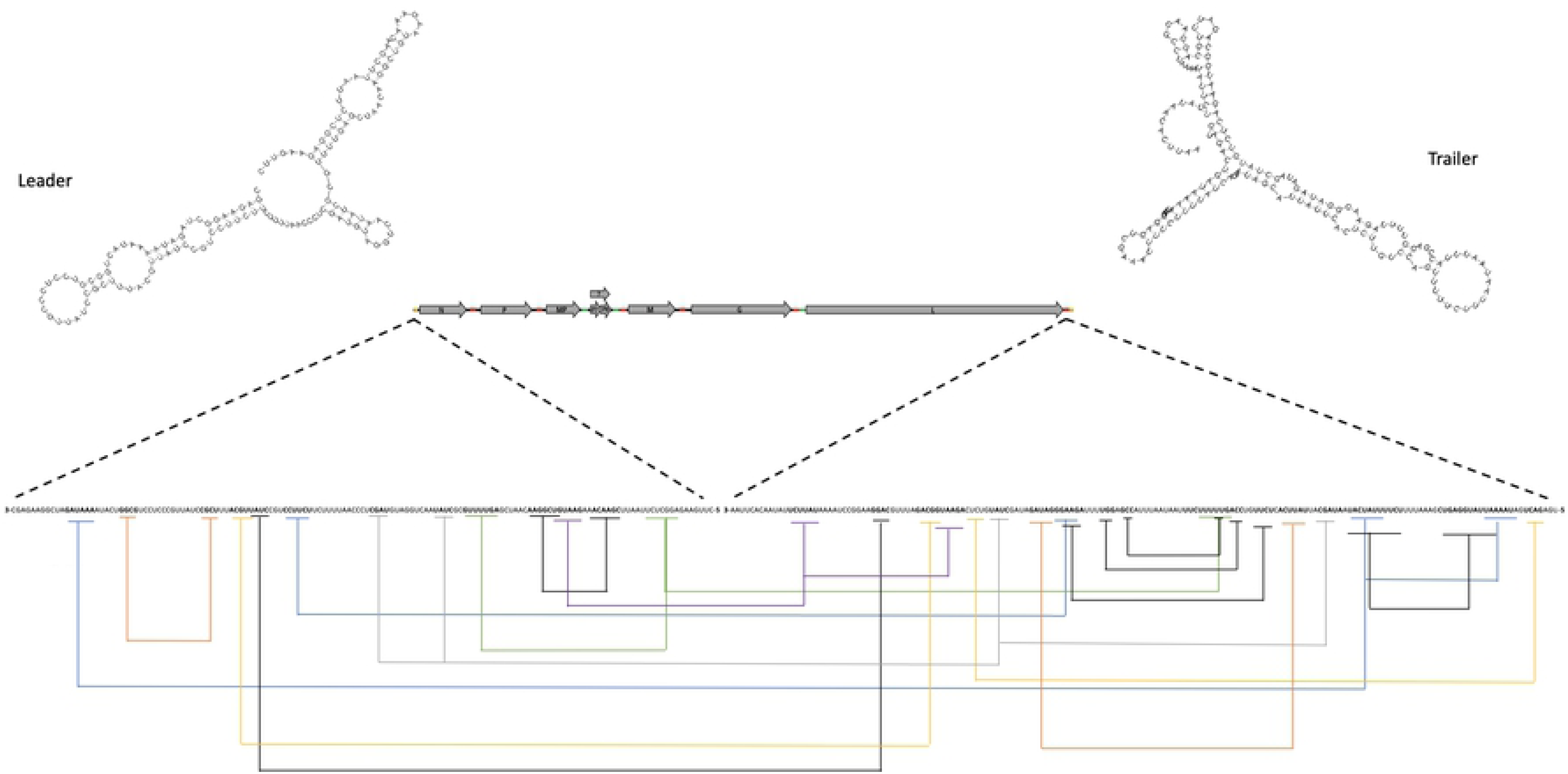
Graphical schematization of terminal regions. Bolded nucleotides denote motifs with complementarity within the same terminal region or between terminal regions. Hypothetical interactions, based on complementarity of each motif, are indicated by coloured solid lines. Predicted secondary structures are shown for both the leader and the trailer.

### Phylogenetic relationships

The multiple sequence alignments for the nucleocapsid (N), glycoprotein (G) and polymerase (L) were 831, 936, and 3,071 amino acids (aa) long, respectively. After eliminating ambiguous fragments, the resulting alignment lengths were 284, 269, and 1,692 aa for each protein, respectively. The best evolution model for N and L was LG+G [34], while WAG+G+I [35] was the best model for the glycoprotein. The two runs for all three proteins and the multi-locus alignment converged and the effective sample size was above 200 for all parameters.

The topology of the multi-locus tree was identical to the polymerase one and congruent with those inferred by analyses of the nucleocapsid and glycoprotein. Cyto- and nucleorhabdoviruses are monophyletic and each genus contains clades with well supported nodes corresponding with their vectors (Fig 3). For maize fine streak virus and rice yellow stunt virus, however, vector-associated phylogenies depended on the protein being analyzed S2 Fig.

**Fig 3.**
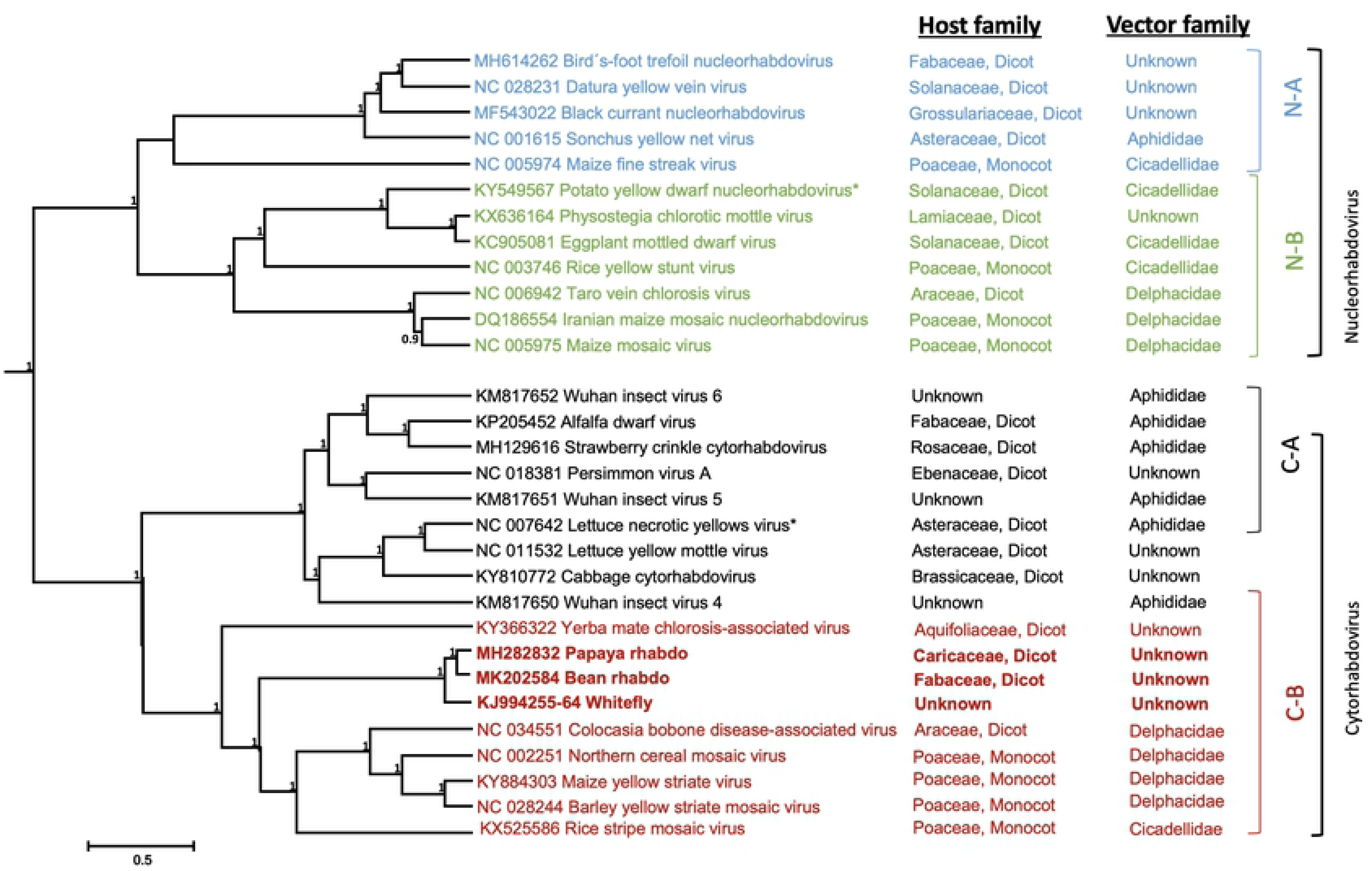
Multilocus phylogeny of plant infecting rhabdoviruses. Multiple sequence alignments of the nucleocapsid, glycoprotein, and polymerase were analyzed in BEAST 1.8.4. Numbers above the nodes represent posterior probabilities, only values above 0.9 are shown. N-A Aphididae transmitted nucleorhabdovirus. N-B Cicadellidae/Delphacidae transmitted nucleorhabdovirus. C-A Aphididae transmitted cytorhabdovirus. N-B Cicadellidae/Delphacidae transmitted cytorhabdovirus. Asterisk denotes type species. The cytorhabdovirus clade is shown in red, where the new papaya virus and its closest relatives are bolded.

The new papaya virus grouped with members of the *Cytorhabdovirus* genus, in a clade that includes the dicot-infecting YmCaV and CbDaV, and the monocot-infecting NCMV, MYSV, BYSV and RSMV, which are transmitted (except for YmCaV) by Delphacidae/Cicadellidae vectors (Fig 3). Interestingly, virus-like sequences allegedly from the *B. tabaci* genome grouped closely with the new papaya virus.

### Genome comparison with a bean-infecting strain

When this manuscript was in final preparation, the genome of a rhabdovirus found in common bean (*Phaseolus vulgaris* L.) was reported from Brazil [36]. Although the sequence record of the papaya cytorhabdovirus reported in the present study has been available since October 1, 2018 (MH282832), it was overlooked and not included in the analysis performed and reported in January 2019 by Alves-Freitas et al [36]. The striking similarities observed between the two genomes and their predicted proteins are summarized in Table 3. The bean cytorhabdovirus has an overall nucleotide identity of 97% with the papaya cytorhabdovirus reported here. This identity level, according to the species demarcation criterium for rhabdovirus species [37], clearly indicates that the bean and the papaya cytorhabdoviruses are strains of the same species. An important difference in the genome organization of the two strains is the presence of hypothetical ORF 4 (nt 3,785 – 4,021) in the bean cytorhabdovirus, which is not present in the papaya cytorhabdovirus.

**Table 3.**
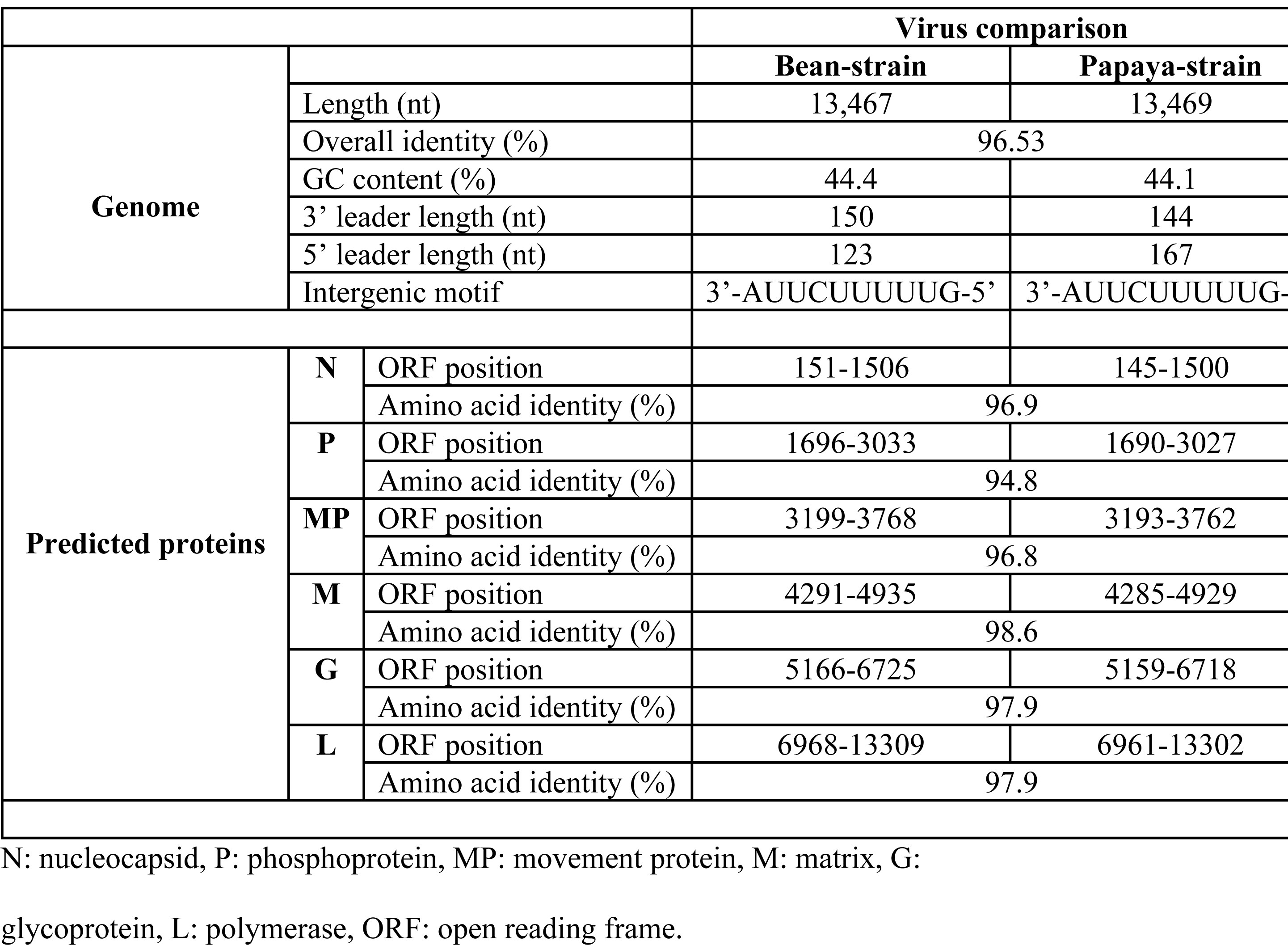
Comparison of genome and predicted proteins between the bean and papaya strains of a new cytorhabdovirus.

### Transmission Electron Microscopy (TEM)

TEM images of the mesophyll cells of papaya leaves infected with PRSV and the new rhabdovirus are shown in Fig 4. Aggregations of rhabdovirus-like particles were detected in the periphery of chloroplasts. Pin-wheel and swirls, as well as crystalline inclusions typical of potyviruses, were readily observed. There were much fewer cells and aggregations of the rhabdovirus-like articles than of potyvirus, which could be related to virus titer. This observation is consistent with the NGS data, where 11.68% of the reads mapped to PRSV and only 0.23% to rhabdovirus.

**Fig 4.**
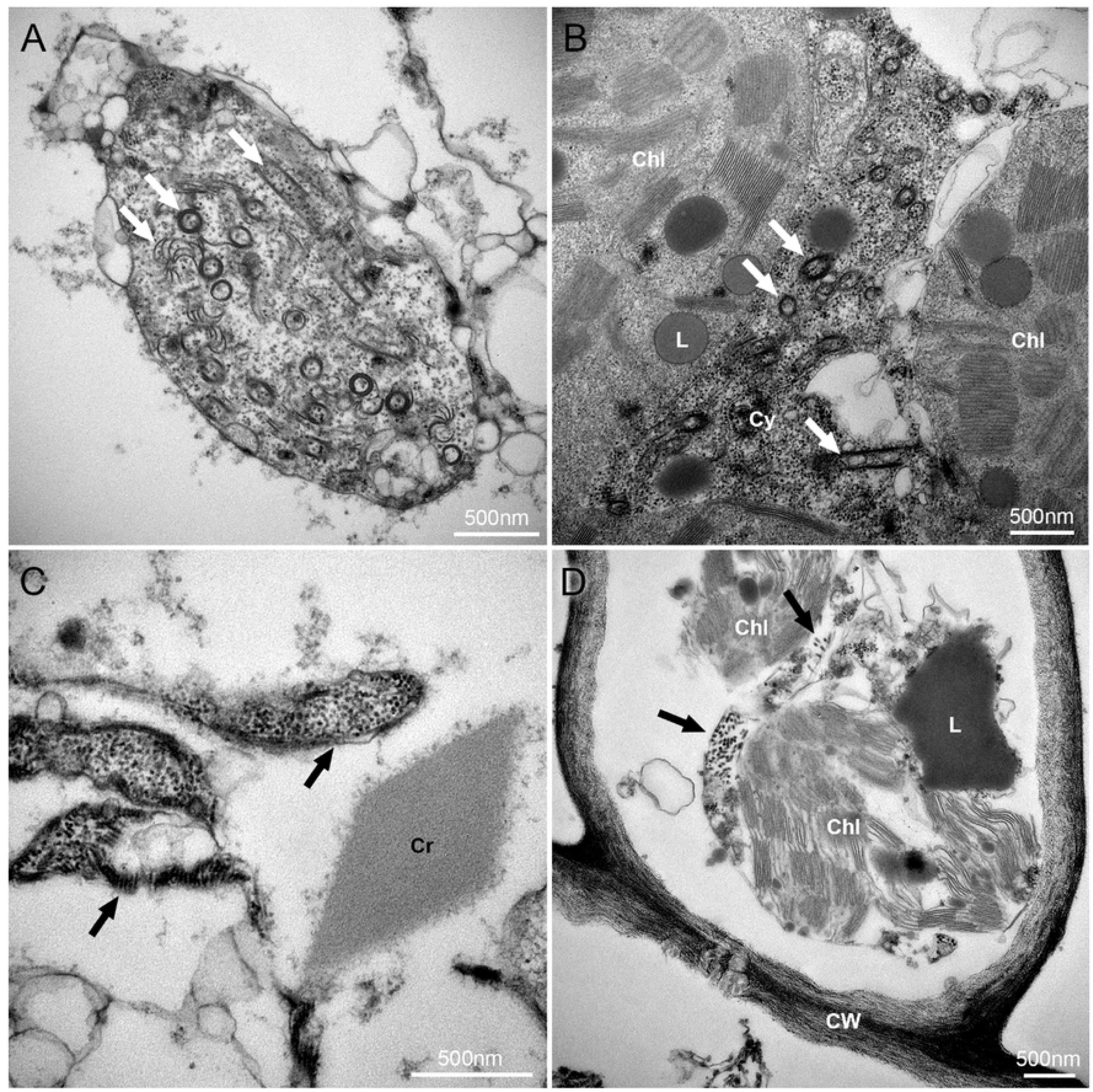
Transmission electron microscopy (TEM) images of the mesophyll cells of papaya leaves infected with papaya ringspot virus and the new cytorhabdovirus. (A-B) White arrows showing potyvirus pin-wheel inclusions in three typical configurations in the cytoplasm. (C-D) Black arrows showing aggregations of rhabdovirus particles in the cytoplasm along with a crystalline inclusion. Chl: Chloroplasts; Cr: Crystalline inclusions; CW: Cell Wall; Cy: Cytoplasm; L: Lipid droplets.

### Virus survey

Primers: (F) 5’-CGCAAAACTCGATTGTTCCG-3’ and (R) 5’ CCTGCTGATGATCCTATCTCC-3’, which amplify a 779 nt region spanning a fraction of the 3’ leader and the nucleocapsid gene, were selected for virus detection. A total of 180 papaya plants in commercial fields from 5 different provinces in Ecuador were tested.

In Los Ríos, the virus was found in 100% (n = 30) of samples (cv. Sunrise) collected from one-year old plants; whereas 13% (n = 30) were positive in an adjacent four-month-old field. In Guayas province, the virus was only detected in one out of 30 plants tested from a two-year-old field. In Manabí, 20% of plants tested positive from a three-year-old field (n = 30). The virus was not detected in selected fields of Sucumbíos, a forest province where ‘Criolla’ papaya was grown, or Santa Elena, where the Hawaiian cultivar Sunset was sampled.

All the plants that tested positive for the papaya cytorhabdovirus were also positive for PRSV. However, no differences in leaf symptoms were observed between PRSV-singly infected plants and plants co-infected with both viruses.

Genome diversity was inferred by comparing an 860 nt fragment of the virus polymerase. Five isolates from Los Ríos, five from Manabí, and only one from Guayas, were selected for RT-PCR using the primers: (F) 5’-GAGAAGTGGAACCTCAATTTCC-3’ and (R) 5’-CTGAAGAGAGAAGGGTCGGT-3’. Sequence alignments of the amplified fragment showed a 99 % identity across isolates from different provinces in Ecuador.

## Discussion

The *Rhabdoviridae* is one of the most diverse virus families as it contains species that infect arthropods, vertebrates and plants [9]. Here, we present the characterization of a new rhabdovirus discovered from papaya plants in Ecuador. Aligning entire genomes of plant rhabdoviruses is difficult due to high divergence of sequences. Nevertheless, the evolutionary history of this virus was confidently inferred using the nucleocapsid, glycoprotein and polymerase amino acid compositions.

The new virus grouped with members of the *Cytorhabdovirus* genus, with rice stripe mosaic virus, yerba mate chlorosis-associated virus and Colocasia bobone disease-associated virus as closest relatives.

In the multi-locus alignment, 75.4 % of the total length corresponded to the polymerase and the resulting multi-locus tree topology was identical to the topology of the polymerase alone, supporting other studies that indicate using the polymerase is an accurate representation of the phylogeny for rhabdoviruses [38].

Furthermore, during this study a strong correlation was revealed between phylogenetically-related species and their vectors. Accordingly, the new papaya rhabdovirus is likely transmitted by a member of the *Delphacidae* or *Cicadellidae*. However, formal transmission experiments are needed to confirm this hypothesis. An interesting finding in this study was the genetic closeness (80 % nucleotide identity) observed between the new papaya virus and sequences from a whitefly genome annotated by Kumar and Upadhyay (unpublished, Genbank acc. numbers: KJ994255-KJ994264). The whiteflies used for the genomic analysis may have been either infected by a cythorhabdovirus or were carrying (potentially as a vector) a cytorhabdovirus acquired from a plant host. These scenarios remain speculative and we have not succeeded in contacting the authors of the whitefly sequence submission.

The genome organization of rhabdoviruses includes five genes flanked by a leader and trailer sequences at the 3’ and 5’ ends, respectively, resulting in the canonical arrangement: 3’-*l*-N-P-M-G-L-*t*-5’.

In plants, cyto- and nucleorhabdoviruses have an additional ORF between the P and M genes, whose product has cell-to-cell movement activity, resulting in the typical 3’- *l*-N-P-MP-M-G-L-*t*-5’ arrangement[13,14]. However, there are numerous examples of divergence from the typical canonical organization for plant rhabdoviruses with additional small ORFs of unknown function [12,39]

In the new papaya cytorhabdovirus, two short ORFs (namely ORFs 7 and 8) were predicted with overlapping termination/initiation codons. This feature is commonly associated with a translation-reinitiation mechanism and has been documented, among others, for some animal rhabdoviruses [5]. The reinitiation mechanism is dependent on TURBS (*termination upstream ribosome binding site*), which includes a pentanucleotide motif that is complementary to the loop region of helix 26 of 18S rRNA [32]. TURBS-like motifs were not identified upstream the reinitiation start codon of ORF 8 in the papaya rhabdovirus. However, the TURBS-reinitiation mechanism has mostly been studied in animal-infecting viruses, and it is possible that a different virus-host interaction, involving the 18S rRNA in plants, operates to promote reinitiation.

One of the genomic hallmarks in rhabdoviruses is the presence of transcription regulatory signals, such as conserved intergenic motifs and self-complementary sequences located at terminal regions [9]. The genome of the new papaya cytorhabdovirus exhibits the conserved motif 3’-AUUCUUUUUG-5’ not only in intergenic regions (except in the MP-P7 junction), but also in the 5’ (*t*) region, suggesting its involvement in transcription termination of the corresponding preceding gene.

In addition, a second conserved motif (here referred to as *intergenic motif II*) was detected in gene junctions MP-P7, P8-M and G-L of the new virus genome (Fig 1), suggesting a potential regulatory role in the transcription of these genes. This motif could not be detected in gene junctions of closely related viruses.

This is the first report of a cytorhabdovirus in papaya and the first sequence deposited for any rhabdovirus of this host. Both its phylogenetic relatedness to cytorhabdoviruses and the TEM observations showing virus accumulation in the cytoplasm support the notion that this virus is not related with the papaya nucleorhabdoviruses reported in the early 1980s [16,17,18]. Since we could not find a papaya plant singly-infected with the new cytorhabdovirus, symptomatology associated to the virus was not determined. Nevertheless, the papaya cytorhabdovirus was detected in field plantings in three different provinces in Ecuador, strongly suggesting this is a naturally occurring virus. Comparisons among the polymerase sequence among 11 of these isolates were highly convergent (99% identity).

Lastly, in January 2019, the genome of a bean-infecting cytorhabdovirus was documented from Brazil [40]. The bean-infecting cytorhabdovirus shares 97% nucleotide identity with the new papaya virus and has similar genome organization (Table 3). Given that the papaya virus genome has been available in Genbank since October 1, 2018 (acc. MH282832) and based on our data that indicates natural field spread, we propose that the bean cytorhabdovirus should be considered a bean-infecting strain of the new species named papaya cytorhabdovirus E. This approach has already been supported by a letter to the editor[40].

## Acknowledgments

The authors thank papaya growers in selected provinces of Ecuador for allowing access to their fields, and Dr. Gary Kinard for critical review. This work was conducted under *Genetic Resource Access Permit* # MAE–DNB–CM–2018–0098 granted by the Department of Biodiversity of the Ecuadorean Ministry of the Environment.

## Supporting information

**S1 Fig. Hypothetical secondary structures predicted for each intergenic región of the new papaya cytorhabdovirus.** Colors indicate probability of complementarity (red: high, green: mild, yellow: low). N: nucleocapsid, P: phosphoprotein, MP: movement protein, M: matrix, G: glycoprotein, L: polymerase, ORF: open reading frame.

**S2 Fig. Single protein phylogenies of the polymerase**, **glycoprotein and nucleocapsid of plant infecting rhabdoviruses.** Bolded accession numbers correspond to species whose evolutionary history is not congruent among proteins. NC_005974 Maize fine streak virus. NC_003746 Rice yellow stunt virus. Numbers above the nodes represent posterior probabilities, only values above 0.9 are shown.

